# Combinatorial action of transcription factors in open chromatin contributes to early cellular heterogeneity and organizer mesendoderm specification

**DOI:** 10.1101/2020.02.26.966168

**Authors:** Ann Rose Bright, Siebe van Genesen, Qingqing Li, Simon J. van Heeringen, Alexia Grasso, Gert Jan C. Veenstra

## Abstract

During gastrulation, mesoderm is induced in pluripotent cells, concomitant with dorsal-ventral patterning and establishing of the dorsal axis. How transcription factors operate within the constraints of chromatin accessibility to mediate these processes is not well-understood. We applied chromatin accessibility and single cell transcriptome analyses to explore the emergence of heterogeneity and underlying gene-regulatory mechanisms during early gastrulation in *Xenopus*. ATAC-sequencing of pluripotent animal cap cells revealed a state of open chromatin of transcriptionally inactive lineage-restricted genes, whereas chromatin accessibility in dorsal marginal zone cells more closely reflected the transcriptional activity of genes. We characterized single cell trajectories in animal cap and dorsal marginal zone in early gastrula embryos, and inferred the activity of transcription factors in single cell clusters by integrating chromatin accessibility and single cell RNA-sequencing. We tested the activity of organizer-expressed transcription factors in mesoderm-competent animal cap cells and found combinatorial effects of these factors on organizer gene expression. In particular the combination of Foxb1 and Eomes induced a gene expression profile that mimicked those observed in head and trunk organizer single cell clusters. In addition, genes induced by Eomes, Otx2 or the Irx3-Otx2 combination, were enriched for promoters with maternally regulated H3K4me3 modifications, whereas promoters selectively induced by Lhx8 were marked more frequently by zygotically controlled H3K4me3. Our results show that combinatorial activity of zygotically expressed transcription factors acts on maternally-regulated accessible chromatin to induce organizer gene expression.

## INTRODUCTION

Cellular heterogeneity increases dramatically during early embryonic development in association with regional specification of the ectoderm, mesoderm and endoderm lineages. During this process, cells respond to extracellular signals as dictated by cell-autonomous constraints such as chromatin state and the presence of factors mediating the response. The embryo is transcriptionally quiescent until zygotic genome activation (ZGA) (Paranjpe and Veenstra, 2015; Vastenhouw et al., 2019), which gradually occurs during the mid-blastula stage in *Xenopus*. This is accompanied by slowing down of cell divisions and introduction of cell cycle gap phases. During the blastula stage, the cells at the animal pole are pluripotent. They are fated to become ectoderm but are competent to respond to mesoderm-inducing signals emanating from vegetal pole cells. During early gastrulation there is a major cellular diversification, concomitant with germ layer formation and morphogenesis.

Within the animal cap at early gastrula stages, ectoderm primarily consists of a superficial (epithelial) layer, and an attached deep (sensorial) layer (Chalmers et al., 2002), which diverge to goblet cells, multiciliated cells, ionocytes and small secretary cells in the epidermal ectoderm by larval stages (Angerilli et al., 2018). In addition, the animal cap cells are competent to form mesoderm, but can also be induced to neural ectoderm by underlying mesoderm during gastrulation. Mesoderm, located at the equatorial region (marginal zone) during early gastrula stages, is induced by nodal-related TGFβ family ligands, produced by vegetal cells. Dorsal mesoderm develops into a signaling center, the Spemann-Mangold Organizer, involved in dorsal-ventral patterning of mesoderm and formation of the dorsal axis during gastrulation (Agius et al., 2000). Using transplantation experiments, it has been established that the dorsal blastopore lip of early gastrula stages induces anterior dorsal structures (head organizer), whereas the blastopore lip at late gastrula stages specifies posterior structures (trunk organizer). Little is known how these activities relate to each other, but it has been shown that inhibition of Wnt signaling is important for the formation of head organizer structures such as anterior endoderm, prechordal plate and anterior chordamesoderm (Niehrs, 1999). The dorsal blastopore lip is also home to superficial mesoderm. Under the influence of Wnt11b and FGF signaling, as well as the Foxj1 transcription factor, involuting superficial mesoderm cells will constitute the ciliated gastrocoel roof that forms the left-right organizer (Schneider et al., 2019; Walentek et al., 2013).

During early development, secreted factors signal to the nucleus of exposed cells, impacting gene expression in conjunction with transcription factors. Chromatin plays a major role in this process, providing the gene-specific permissive or restrictive context for transcription (Jambhekar et al., 2019; Perino and Veenstra, 2016). This involves histone modifications such as the permissive promoter mark H3K4me3 and the repressive Polycomb mark H3K27me3, both of which increase dramatically during early *Xenopus* development (Bogdanović et al., 2012; Hontelez et al., 2015a; Paranjpe and Veenstra, 2015). A large majority of genomic loci is decorated with these histone modifications by maternal factors; these loci are collectively referred to as the maternal regulatory space, whereas a relatively small number of promoters requires new embryonic transcription for the acquisition of H3K4me3 or H3K27me3 (Hontelez et al., 2015b).

Very little is known how cellular heterogeneity and the cellular responses to extracellular signaling relate to chromatin accessibility and how this is regulated regionally during early development. Here we report on the regulation of chromatin accessibility during early development and in early gastrula animal cap and dorsal marginal zone using ATAC-sequencing. The regional differences in gene expression and chromatin accessibility were related to cellular heterogeneity observed in whole embryo single cell RNA-sequencing (scRNA-seq) and scRNA-seq data of dissected animal cap and dorsal marginal zone tissue. We inferred transcription factor activities in specific cell clusters, which were tested in animal caps by microinjection and RNA-sequencing. We assessed the extent to which organizer-expressed transcription factors activate gene expression within open chromatin, and how this relates to H3K4me3 promoter marking by maternal factors. The data support the early emergence of head and trunk organizer as well as superficial mesoderm cells. Moreover, the data show how early cellular heterogeneity emerges in response to inductive events by action of zygotic transcription factors in the context of maternally marked, accessible promoters.

## RESULTS

### Dynamics of chromatin accessibility during early development

To define the chromatin regulatory landscape during early development, we performed ATAC-seq of biological replicates for blastula, early and late gastrula, and neurula stages (respectively stage 9, 10½, 12, and 16; Figure 1A). We clustered the open chromatin ATAC-seq peaks together with H3K4me3 and p300 ChIP-seq data (Hontelez et al., 2015a) of the same developmental stages. Open chromatin as observed by ATAC-seq is found in regions with H3K4me3 (promoters), the co-activator p300 (enhancers) or both (promoter-proximal regulatory elements) (Figure 1A). H3K4me3-decorated promoter regions displayed higher levels of chromatin accessibility than enhancer elements that recruit p300 (Supplemental Fig. S1A). Generally, ATAC-seq enrichment increased from stage 9 onwards, both for H3K4me3-positive promoters and p300-bound enhancers (Supplemental Fig. S1A). Before stage 9, we have not been able to obtain high-quality ATAC-seq tracks because of a lack of enrichment, which may suggest that regulatory elements become only accessible after the mid-blastula transition.

**Figure 1.**
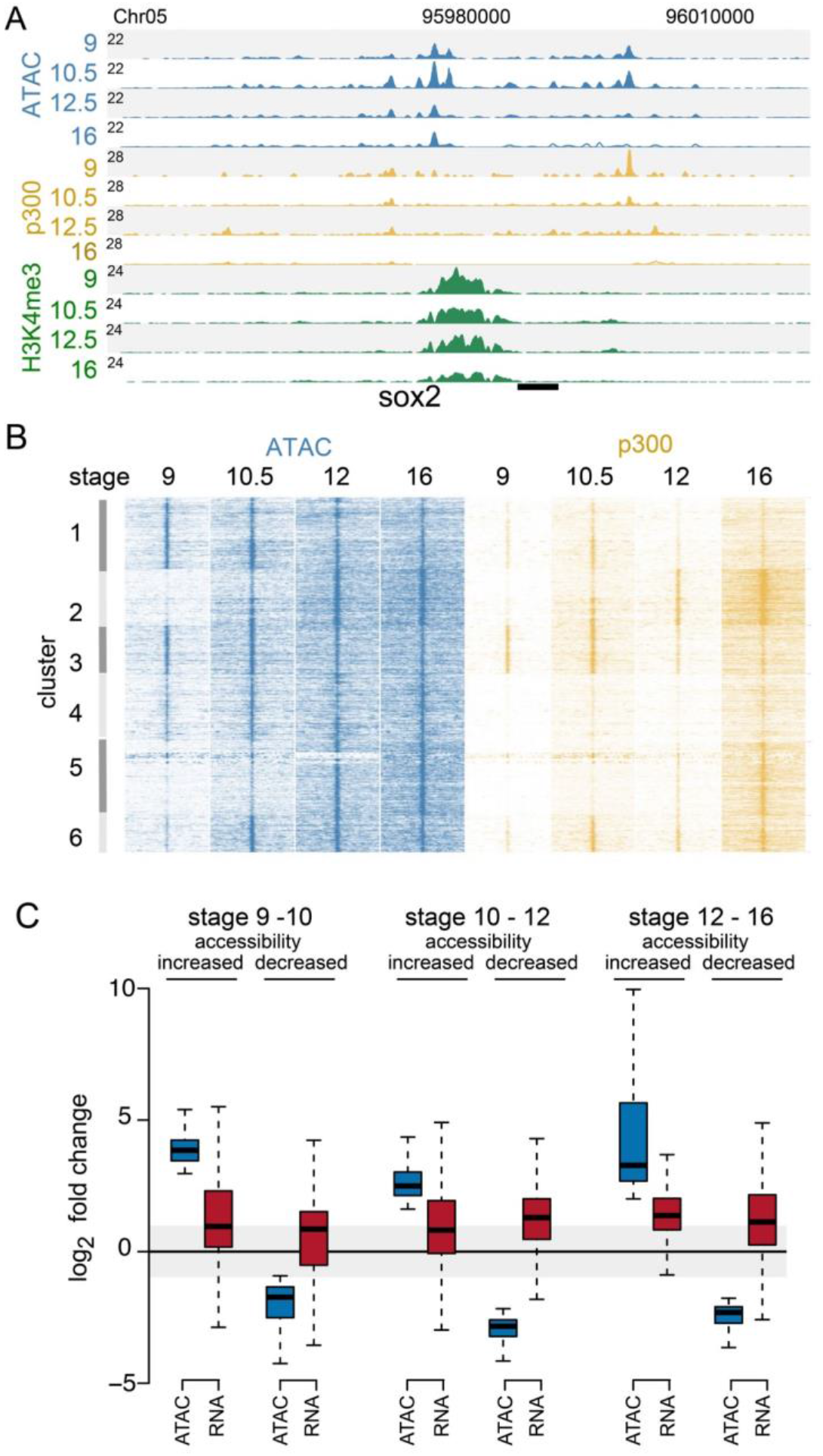
Chromatin accessibility during early development. **A.** Genome browser view showing chromatin accessibility (ATAC-seq) and ChIP-seq (p300 and H3K4me3) profiles at stage 9, 10½, 12 and 16 around the *sox2* locus. ATAC-seq peaks were found at promoter (H3K4me3) and enhancer (p300-bound) regulatory regions. **B.** Chromatin accessibility and p300 binding at differential ATAC-seq peaks visualized using K-means clustering. **C.** Box plots showing sequential-stage comparisons of fold change in accessibility (ATAC-seq) and corresponding changes in gene expression (RNA-seq).

Pair-wise comparison of sequential stages yielded over 7000 differentially accessible regions (see Methods) (Figure 1B). Clustering of these regions with p300 and H3K4me3 data revealed that the majority of these regions with dynamic open chromatin, are enriched for p300 but not H3K4me3 (Supplemental Figure S1B). Moreover, the ATAC-seq and p300 signal intensities correlate for these dynamic open chromatin regions, suggesting they represent developmental stage-specific accessible enhancers. Clusters 2, 4, 5, and 6 showed accessibility signals increasing from stage 9 to stage 16, whereas cluster 1 and 3 consisted of regions showing a reduction in the signal after gastrulation. To assess the extent to which the open chromatin dynamics are linked to gene regulation, we analyzed the transcript levels of nearby genes. We found for each of the sequential stage comparisons that genomic elements with increased chromatin accessibility are associated with an increase in expression of the associated genes (Figure 1C; left side of each panel). Surprisingly, genes associated with individual genomic elements showing a decreased accessibility also upregulated in many cases. It should be noted that many of these genes are regulated by multiple enhancers, sometimes with different dynamics. In addition, transcript stability may cause transcript dynamics to lag behind chromatin accessibility dynamics. The results raise the question of how changes in chromatin accessibility relate to regional specification, germ layer formation and the heterogeneity in gene expression programs associated with the onset of gastrulation.

### Mesoderm-induced genes exhibit open chromatin in animal caps

To determine the extent to which chromatin accessibility is spatially acquired in early gastrula stage embryos (stage 10½), we preformed ATAC-seq on ectodermal (animal cap: AC) and organizer (dorsal marginal zone: DMZ) explants (Figure 2A). Ectoderm-expressed genes such as *tfap2a* and *grhl3* showed high accessibility at regulatory regions in the AC and relatively low signals in DMZ (Figure 2A). By contrast, the organizer-expressed genes *gsc* and *chrd* were equally accessible in AC and DMZ explants. To assess how general these observations are, we performed differential gene expression analysis (fold change >= 2 and FDR < 0.05) of AC and DMZ RNA-seq samples (Blitz et al., 2016). Differential genes were then linked to their closest ATAC-seq peak to see how well spatial expression differences match with differences in chromatin accessibility (Figure 2B). We observed that genes with higher expression in AC compared to DMZ, also exhibit higher chromatin accessibility in AC. However, for genes with higher expression in DMZ, the associated regulatory regions showed a similar accessibility signal in both explants, similar to what was observed at *gsc* and *chrd*. AC cells are considered pluripotent at the blastula stages and lose competence for mesoderm induction during gastrulation (Borchers and Pieler, 2010; Jones and Woodland, 1987). Consistent with the competence for mesoderm induction, mesoderm-expressed genes appear to exhibit accessible chromatin in AC.

**Figure 2.**
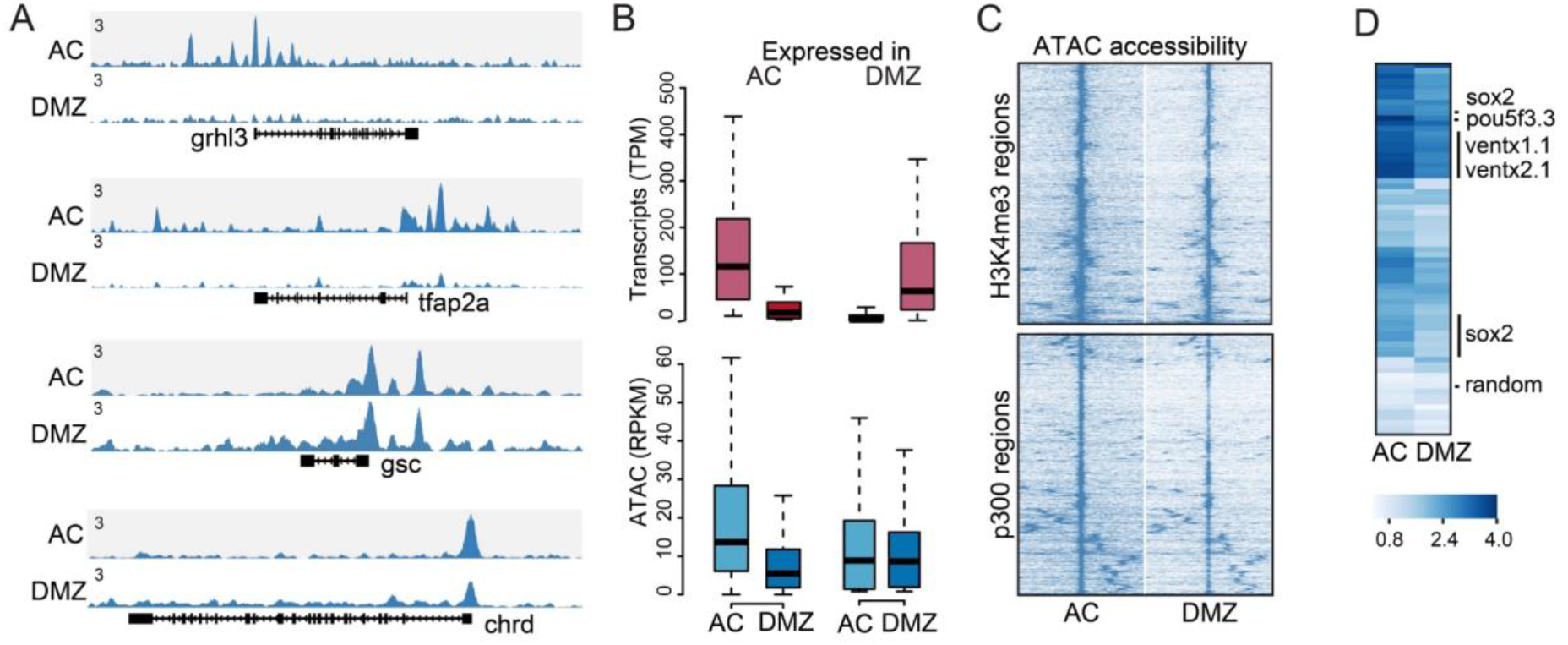
Chromatin accessibility in animal cap (AC) and dorsal marginal zone (DMZ). **A.** Genome browser view of AC and DMZ accessibility profiles for ectoderm-expressed (*tfap2a* and *grhl3*) and organizer-expressed (*gsc* and *chrd*) marker genes. **B.** Boxplots showing differential gene expression (AC versus DMZ) and associated ATAC-seq signals. **C.** Hierarchical clustering of AC and DMZ ATAC-seq data on H3K4me3-positive (top) and p300-positive (bottom) ATAC-seq peaks. **D.** Heat map showing accessibility signal at p300-positive ATAC-seq peaks surrounding pluripotency genes. The row labeled ‘random’ shows accessibility signals at random genomic loci.

To further characterize regional chromatin accessibility at promoters and enhancers, we divided AC and DMZ peaks into p300 and H3K4me3-positive regions. Hierarchical clustering showed a somewhat lower ATAC signals in DMZ for both H3K4me3 and p300 positive regions (Figure 2C), but the significance of this result is not clear. Most of the p300-associated regulatory elements of pluripotency genes (*pou5f3.3, pou5f3.1, sox2*, and the *Nanog*-related genes *ventx2.1*, and *ventx1.1* (Scerbo et al., 2012) were accessible in both AC and DMZ, although to variable degrees, whereas random genomic regions were not accessible in either tissue (Figure 2D). The transcription start sites of these pluripotency genes also showed relatively strong chromatin accessibility in AC compared to DMZ (Supplementary Figure 2A). Overall these observations confirmed that regulatory regions in AC cells have relatively open chromatin, irrespective of the transcriptional activity of the associated genes, in line with the pluripotent nature of the AC and its and competence for mesoderm induction.

We wondered whether we could identify regulatory factors driving differential chromatin accessibility between AC and DMZ by integrating ATAC-seq and RNA-seq data. We used Gimme Motifs maelstrom (Bruse and Heeringen, 2018), an ensemble regression algorithm, to identify motif activity associated with differential chromatin accessibility between AC and DMZ. We plotted the differential expression of the transcription factors capable of binding these motifs together with the inferred motif activity (Supplementary Figure S2B-C). The differentially expressed transcription factor genes with chromatin accessibility-associated motifs in AC included factors involved in ectodermal and epidermal development (*tafp2a, tfap2c, grhl1*) (Luo et al., 2005; Tao et al., 2005). In addition, several Klf genes (*klf2, klf5, klf17*) were identified which are highly expressed in the animal cap cells of *Xenopus* and are involved in pluripotency and self-renewal in mammalian cells (Gao et al., 2015). For the DMZ, differentially expressed genes with accessibility-associated DNA-binding motifs included *eomes*, and Forkhead factors such as *foxa1* and *foxd3*, which are known to play roles in mesendoderm specification and axial mesoderm (Charney et al., 2017; Steiner et al., 2006). These results highlight the regional and differential activities of these factors in the context of broadly accessible chromatin in the early gastrula embryo.

### Single-cell analysis of spatiotemporal trajectories of ectodermal and mesendodermal cell states

Our results indicated that chromatin accessibility differs regionally in the early embryo for some genes, but not for others. During gastrulation, cellular heterogeneity is known to increase dramatically at the transcriptional level. For example, single cell profiles sampled from blastula to tailbud stage embryos (stages 8-22) has documented the emergence of an increasing number of cell states during early development (Briggs et al., 2018). We analyzed stage 8, 10, and 12 whole embryo single-cell data as a first step to assess the emerging heterogeneity during gastrulation and the associated developmental cell trajectories. We filtered, normalized and visualized the data with a force-directed layout, using available stage and cell type annotations (Figs. 3A, S3A). We called Louvain clusters (cell clusters L0-L20) and hypervariable genes (differential between cell clusters). We labeled cell clusters based on predominant cell annotations in these clusters (Supplemental Fig. S3A-B). The cells annotated as stage 10 neural ectoderm express both *sox2* and *tfap2a*, whereas *tfap2a* is not expressed in neural ectoderm at later stages (Fig. S4). This suggests that these cells are closely related to non-neural ectoderm at early gastrula stages, consistent with Louvain clusters of cells with mixed neural and non-neural annotations at this stage (clusters L3, L16, L18, L19, Supplemental Fig. S3A-B). To assist the interpretation, we color-coded clusters based on similarity in cell type annotations and gene expression (Figs. 3B, S3B-C, S4). Stage 8 blastomeres are relatively homogeneous, represented by a single cluster (L4) that is most related to clusters comprised of non-neural/neural ectoderm at stage 10 (L3, L8, L18; Figs. 3B, S3B). From these basal stage 10 clusters, there is a continuous trajectory to stage 10 clusters with neural ectoderm annotation (L5), mixed neural ectoderm and marginal zone annotations (L10), marginal zone (L12) and organizer mesendoderm (L2). Cells in stage 10 cluster L10, with mixed marginal zone and neural ectoderm annotations, express both *t (tbxt)* and *sox2* in the same cells, but low levels of *tfap2a*, in line with a potential bi-potent neuro-mesodermal cell state (Supplemental Figs. S3B, S4). Endoderm cells are rather sparse in this data set, but an ectodermal trajectory from basal ectoderm clusters to stage 10 non-neural ectoderm can be observed. Clusters comprised mainly of stage 12 cells are more located to the periphery relative to the stage 10 clusters, with a distinct stage 12 neural ectoderm cluster that is juxtaposed to stage 10 ectoderm (L18) and neural ectoderm (L5). Stage 12 involuted dorsal mesoderm (L9) is juxtaposed to stage 10 organizer (L2) and marginal zone (L12), as well as a stage 12 cluster with tailbud cell annotations (L12; Figs. 3B; S3B-C).

**Figure 3.**
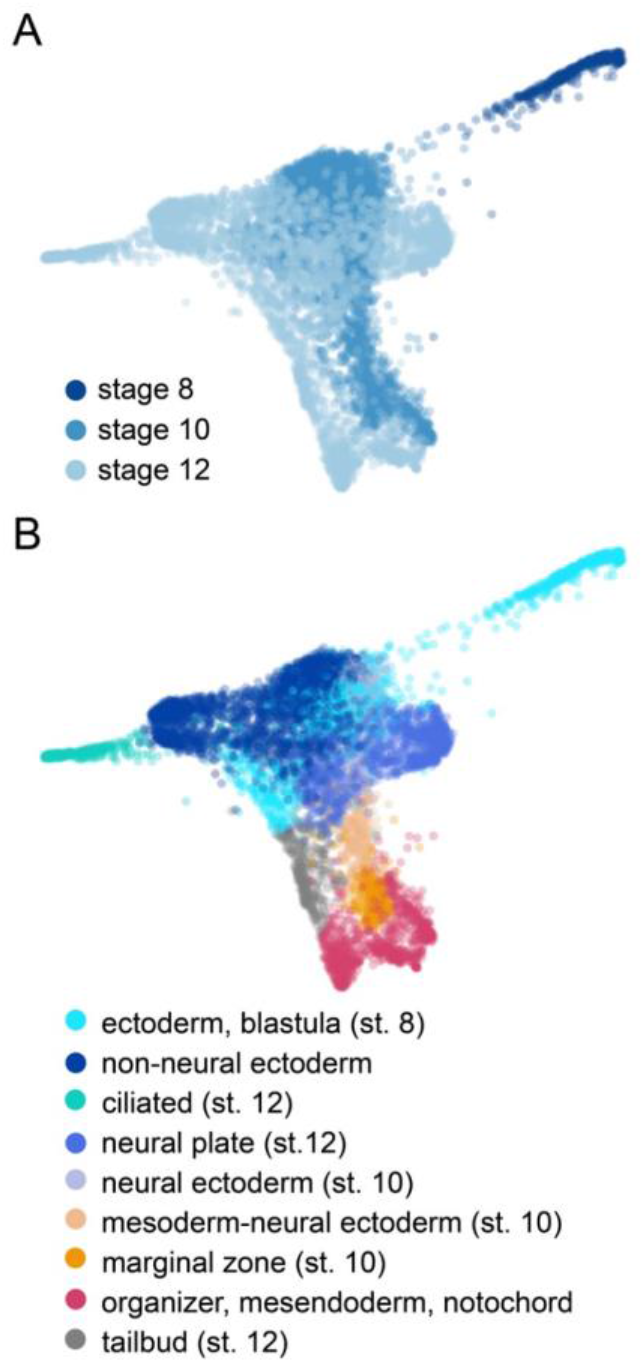
Cellular heterogeneity and developmental trajectories in blastula and gastrula stages. Force-directed layout of whole embryo scRNA-seq for stages 8,10 and 12 colored by stage (**A**) and cell type annotations (**B**).

These whole embryo single cell profiles lack spatial information, although the localization of cell clusters can be tentatively inferred from highly expressed genes. To define the early events associated with the specification of mesodermal and epidermal lineage during gastrulation, and to link single cell transcriptomic profiles to spatially localized gene regulation, we performed single cell RNA-sequencing on dissected animal cap (AC) and dorsal marginal zone (DMZ) explants. We analyzed hand-picked cells from dissociated stage 10½ AC and DMZ explants, collected from two different experiments (Methods). We obtained seven single cell clusters (C0-C6), with the cells of the same region of the embryo generally clustering together (Fig. 4A-B). Based on known marker gene expression of *sox11* (ectoderm inner layer) and *grhl3*, *krt*, and *upk3b* (ectoderm outer layer) (Chalmers et al., 2006), we tentatively assigned AC clusters C0 and C1 to these ectodermal layers (Figs. 4C-D, S5A-B, S6). Both clusters express relatively high levels of the pluripotency factor-encoding transcripts *pou5f3.3* and *sox2*. These clusters also express *ventx1.1* and *ventx1.2*, the closest amphibian homologs of the mammalian Nanog protein (Scerbo et al., 2012), which are abundant in the ventral and animal cap regions (Supplementary Figs. S5, S7). Clusters C2, C3, and C4 all express high levels of mesendodermal markers such as *t* (*tbxt*), *vegt*, and *mix1*. C2 shows the highest levels of *wnt11b* and early expression of *foxj1* (Walentek et al., 2013), which mark respectively involuting mesoderm and superficial mesoderm, the epithelial layer of involuting mesoderm. C3 expresses the highest levels of well-known organizer genes such as *gsc*, *otx2* and *chrd*, in addition to endodermal markers such as *gata4*, *sox17a and sox17b* (Fig. 4C, S5-6). In addition, C3 cells express the head organizer genes *cer1*, *dkk1*, *frzb*, and *fst*, suggesting these cells comprise the precursors of anterior endoderm, prechordal plate mesendoderm and anterior chordamesoderm. C4 expresses also organizer genes such as *gsc*, *otx2*, and *chrd*, but compared to C3, C4 cells express lower levels of head organizer genes and relatively high levels of *cdx4*, *t* (*tbxt*) and *irx3* (Figs. 4C-D, S5-7). This cluster likely constitutes non-involuted mesoderm and prospective trunk organizer cells. C5 is derived from the AC and shows both inner layer characteristics (*sox11*) and expression of *foxj1* and *klf5* (Fig. S5), therefore most likely constituting progenitors of ciliated cells. C6 exhibits a mosaic of ectodermal (*sox11, sox2*) and mesodermal expression (*t, chrd, eomes, vegt*) that is present at the level of individual single cells (Fig. 4D, S6). Genes specifically expressed in this cluster, such as *fgf8* and *foxb1*, are expressed in the upper blastopore lip (Fig. S7), in the inner layer of dorsal ectoderm and in non-involuted mesoderm (Gamse and Sive, 2001; Gentsch et al., 2013; Pera et al., 2014; Takebayashi-Suzuki et al., 2011).

**Figure 4.**
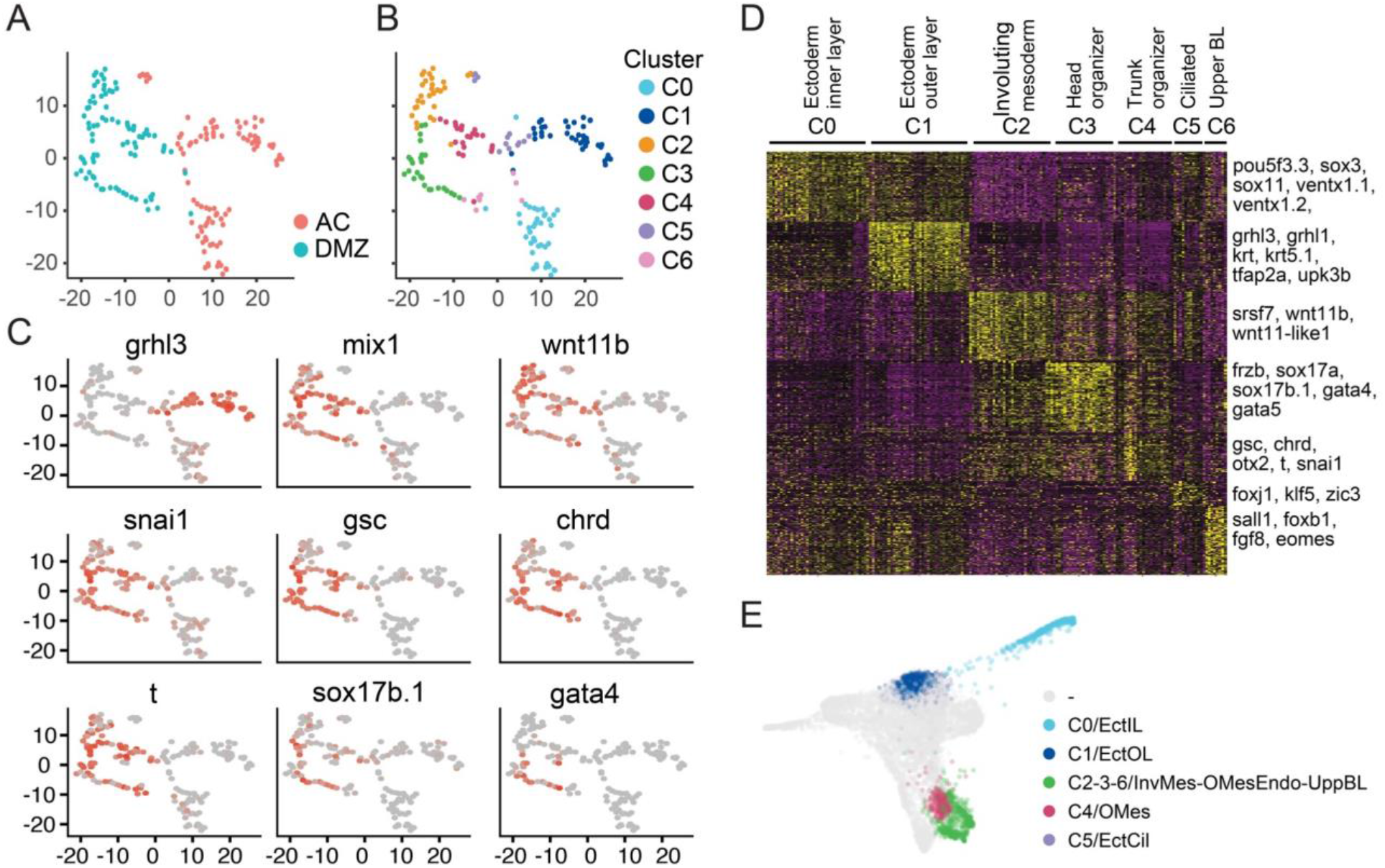
scRNA-seq of hand-picked cells from stage 10½ dissected AC and DMZ explants. **A.** Dimensionality reduction using t-SNE. Each dot represents a cell. Colors indicate AC and DMZ. **B.** As panel A, colors indicate clusters (C0-C6). **C.** Feature maps, showing the expression of selected genes in single cells (cf. Supplemental Fig. S5). Color scale represents the gene expression value for each cell for a given gene, from low (grey) to high (orange). **D.** Heat map depicting top 100 hypervariable genes in each cluster (cf. Supplemental Fig. S6 for high resolution version with all gene names). **E.** Spatially restricted AC and DMZ cell clusters embedded in force-directed graph of whole embryo scRNA-seq data (cf. Fig. 3).

To place our spatially localized AC and DMZ single cell clusters in the whole embryo stage 8-12 dataset, we determined the correlations of the AC and DMZ clusters (C0-C6) with all clusters in the whole embryo data set (L0-L20). As expected, AC cluster C0, C1 and C5 correlated mostly with cells annotated as stage 10 (neural / non-neural) ectoderm and stage 8 blastomeres (Fig. 4E, S8). Involuting mesoderm (C2) and head organizer mesendoderm (C3) correlated best with cells annotated as organizer (L2), with C3 matching L2 best. Organizer mesoderm cluster C4 correlated with both organizer and marginal zone (L12, L2), whereas upper blastopore lip cluster C6 correlated only weakly with stage 10 organizer (L2). Overall, these combined analyses not only confirm known cellular ontologies, they also chart trajectories of the cellular heterogeneity that arises during gastrulation in the AC and DMZ. The data also highlight the progressive nature of germ layer specification, with partially overlapping cell states across developmental stages.

### Integration of single cell transcriptome clusters and chromatin accessibility

To identify potential regulators of cluster-specific gene expression, we analyzed the regulatory elements associated with the hypervariable genes, i.e. transcription factor motifs in regulatory elements near genes with cluster-enriched gene expression. Similar to the analysis of chromatin accessibility in AC versus DMZ, we used GimmeMotifs maelstrom (Bruse and Heeringen, 2018), but now regressing motifs in open chromatin to the variance in gene expression between single cell clusters. The ‘motif activity’ in this context represents the relation between the motif score in accessible regions and the variance in gene expression associated with these elements (Fig. 5A).

**Figure 5.**
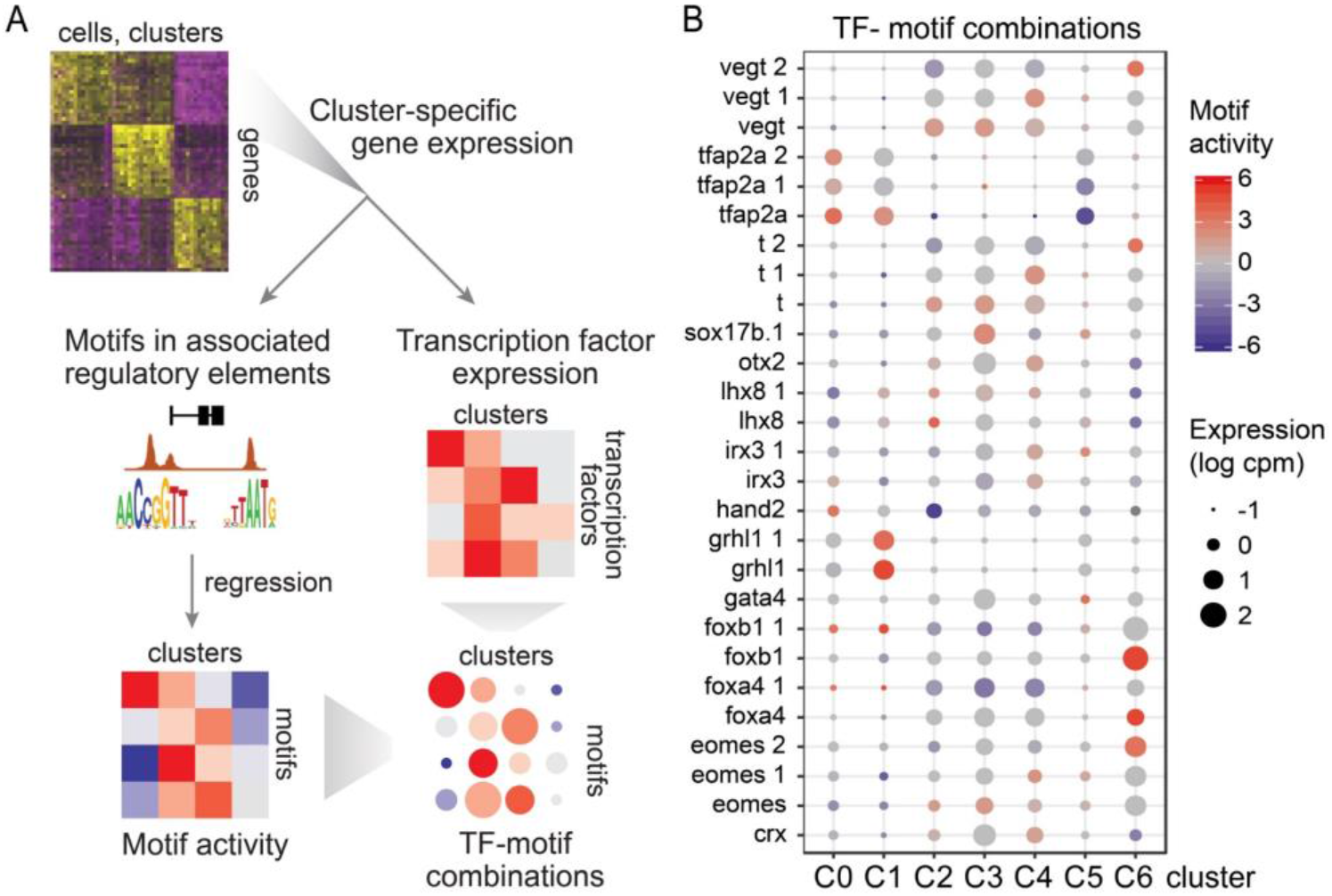
Integration of single transcriptomics and chromatin accessibility, for identifying regulators driving cluster specific gene expression. **A.** Schematic overview, outlining the steps involved in the integrative analysis: (1) Identifying transcription factor motifs associated with regulatory regions closest to cluster-specific genes, and regression to cluster gene expression. 2) Prioritizing transcription factors based on their gene expression and motif activity in specific clusters. Combining the information (lower right panel) on motif activity (color) and corresponding transcription factor expression allows prediction of factors that may play a role in cell cluster-specific gene expression. **B.** Heat map of transcription factor-motif combinations showing cluster-specific motif activity (z-score, color) and gene expression (size of dot).

In the next step, we determined which TFs are predicted to bind these motifs, prioritizing transcription factors based on both their expression and corresponding motif activity in cell clusters (Fig. 5A-B). This analysis is agnostic to whether DNA binding proteins activate or repress transcription, as both positive and negative contributions to cluster gene expression are included. A positive motif activity in a particular cluster means that the motif can explain part of the variance in hypervariable gene expression; genes with the motif tend to be more abundantly expressed in that particular cluster. The transcription factor-motif combinations uncovered in this way include many known well-known regulators, for example Tfap2a (AC clusters C0 and C1, but not C5), Grhl1 (C1), Vegt and T (DMZ clusters C2-4, C6, through different motifs), and Otx2 and Crx (C4) (Fig. 5B). Less well-known is Foxb1; both motif activity and cluster-enriched expression support a potential role in upper blastopore cluster C6, similar to Eomes. Irx3 is expressed in organizer cluster C4 with associated positive motif activity. Lhx8 expression is relatively low at stage 10½ and but its motif activity was mostly restricted to C2 cells, with some activity in C3 and C4 where Lhx8 is also expressed. These results identify potential regulators of the gene regulatory network in early gastrula embryos. This raises the question how these transcription factors contribute to gene regulation, and whether some of these factors can act in a combinatorial fashion in promoting cluster-specific or regional gene expression.

### Combinatorial action of TFs induces organizer gene expression in animal cap explants

To address the contributions of the transcription factors identified by integrated analysis of single cell transcriptomes and chromatin accessibility, we selected candidates for functional characterization based on cluster-specific motif activity and gene expression in DMZ clusters. We selected Lhx8 (C2-C4), Irx3 (C4), Otx2 (C4), Foxb1 (C6) and Eomes (C6) to test their ability to induce DMZ gene expression in AC. We injected synthetic mRNA encoding these factors in the animal pole of one-cell stage embryos, cut animal caps at the blastula stage (stage 8), and collected the animal caps for RNA-sequencing when control embryos reached stage 10½. We also injected combinations of *foxb1* and *eomes*, as well as *irx3* and *otx2* mRNA. Differential gene expression analysis identified a total of around 600 genes (Figs. 6A, S9; Supplemental Table S1) that were differentially expressed (DE) in the overexpressing AC explants compared to water-injected AC explants (adjusted p-value < 0.05, fold change >= 2). Among individually overexpressed transcription factors, *eomes* and *otx2* overexpression had a larger effect on the transcriptome compared to *foxb1, irx3* or *lhx8* overexpression (Figs. 6A, S9). The combination of *foxb1* and *eomes* resulted in a marked difference in transcriptional response, with some genes activated more strongly and other genes less strongly compared to *eomes* alone. For example, whereas *fst* was robustly upregulated in *eomes* but not in *eomes-foxb1*-injected caps, *frzb* was induced only in *eomes-foxb1* caps. The combination of *irx3* and *otx2* resulted in a pattern of up- and downregulation that was similar to that caused by *otx2* alone, but a relatively small number of genes was upregulated more strongly when the two factors were combined. *Lhx8*-injected caps showed less profound activation of DMZ-expressed genes compared to *eomes* or *otx2* injections. Nonetheless, several of the most strongly induced genes in *foxb1-eomes* caps were also expressed in *lhx8* caps. In all these AC explants with overexpressed DMZ factors, genes with mesendodermal or organizer expression such as *t (tbxt), mixer, vegt, foxa1, nog, foxd4, chrd, gsc*, and *sox17* were activated, whereas the expression of AC genes like *tfap2a* and *foxi4.2* was reduced (Figs. 6A, S9).

**Figure 6.**
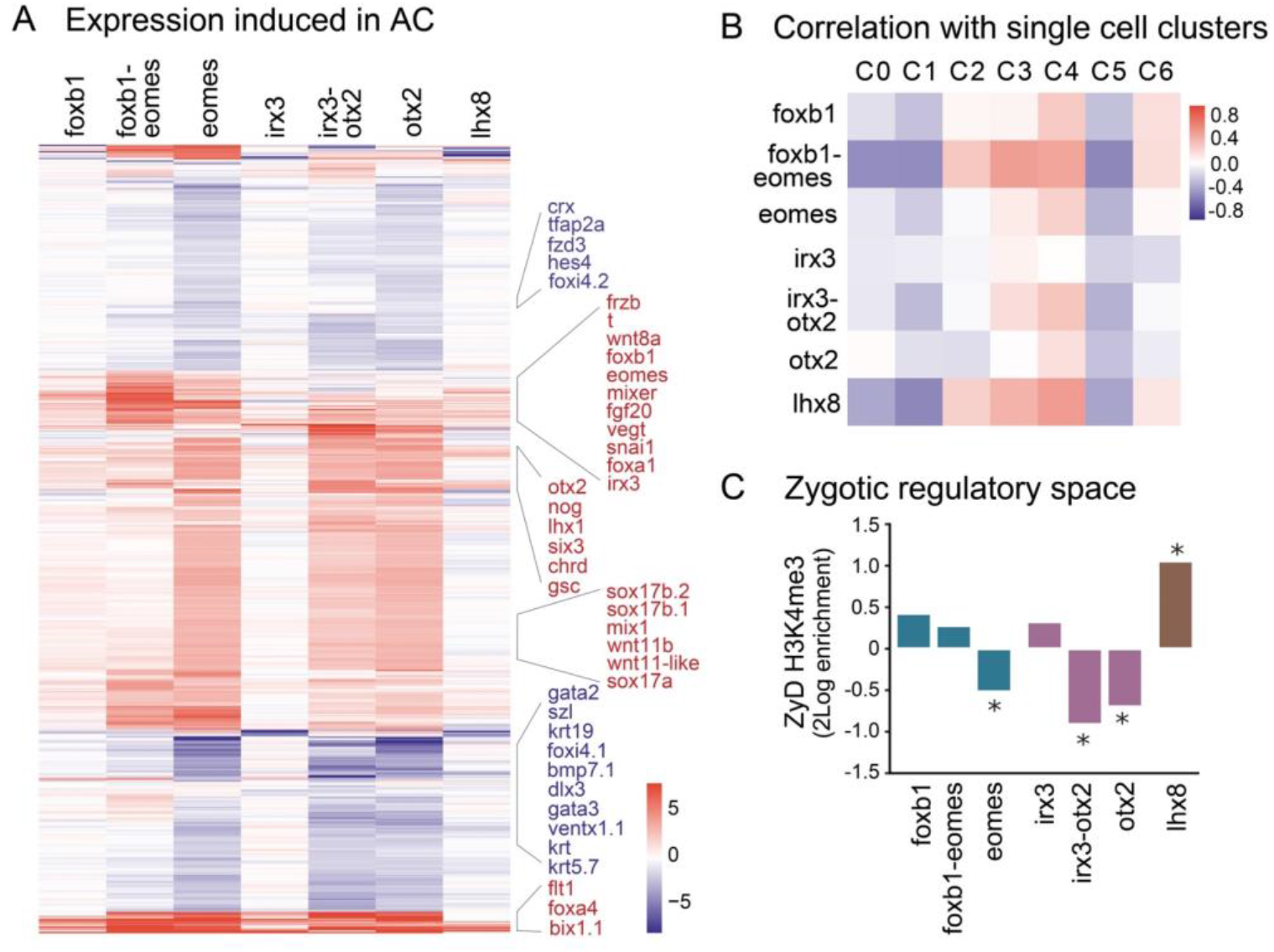
Induction of organizer gene expression in AC cells. **A.** Heat map showing log2 fold expression changes of differentially expressed genes in AC tissues overexpressing Foxb1, Foxb1-Eomes, Eomes, Irx3, Irx3-Otx2, Otx2 and Lhx8. **B.** Correlation heat map of overexpression RNA-seq samples and single cell clusters. **C.** Fold enrichment of genes with zygotically defined (ZyD) H3K4me3 at their promoter in AC with transcription factor overexpression. Asterisk indicates hypergeometric p-value <= 0.01.

To define the induced cell states, we compared the profiles of overexpression RNA-seq samples and the single cell clusters. We performed a correlation analysis using the expression values of genes common between differentially expressed genes in the bulk data set, and hypervariable genes in the single cell data set (Figure 6B). Eomes by itself induced transcriptomic changes that are related to the gene expression profiles of C3-C4 cells (organizer), and Foxb1-induced transcripts correlated with C4 (presumptive trunk organizer) and C6 (upper dorsal blastopore lip) expression. In combination, however, these factors increase the transcriptome similarity to C2-C3-C4 cells, whereas the correlation with C6 gene expression is similar to that caused by Foxb1 alone. Irx3 and Otx2 individually cause some similarity to respectively C3 and C4 cells, whereas in combination the correlation with both C3 and C4 cells is increased. Lhx8 overexpression caused expression of organizer and mesodermal genes (*gsc, chrd, t, mespb, wnt11-like*) and the induced transcriptome correlated most strongly with C4, with lower correlations with C3 and C2 cells.

These data show that all these factors induce transcriptional changes related to mesodermal and organizer cells, and moreover that Foxb1-Eomes and Irx3-Otx2 cause qualitatively and quantitatively different effects when expressed in combination. Notably these effects differ for specific spatial gene expression programs, as Foxb1-expressing AC cells partially recapitulate the transcriptome observed in upper blastopore lip (C6) cells, but the resemblance to C6 expression is unaffected by Eomes co-expression.

Previously, we defined sets of promoters based on whether they required embryonic transcription for gaining the permissive promoter mark H3K4me3 (Hontelez et al., 2015a). Maternally and zygotically defined (MaD, ZyD) H3K4me3 on promoters is related to DNA methylation; unmethylated CpG island promoters acquire H3K4me3 independent of embryonic transcription, revealing mechanistically different modes of transcriptional activation. We wondered to what extent the transcription factors tested in this study, could activate genes in zygotic regulatory space, alone or in combination. We tested if genes more than two-fold upregulated with these transcription factors were enriched for ZyD H3K4me3 genes. We found insignificant or no enrichment of ZyD genes among genes activated by individual transcription factors in AC, with the exception of Lhx8 (hypergeometric p-value 0.001; Fig. 6C). Rather, ZyD H3K4me3 genes were depleted among genes which expression was increased in Eomes (p-value 0.006), Otx2 (p-value 0.0009) and Irx3 plus Otx2-injected (p-value 0.00004) animal caps. We also wondered if the Foxb1-Eomes and Irx3-Otx2 combinations could activate genes synergistically and identified just over 40 genes with more than two-fold cooperativity for each group (fold change >=2 and fold change >= 2x the product of individual fold changes). *Frzb* and *chrd* were among the top genes induced synergistically by Foxb1-Eomes, whereas and *dmbx1 (otx3), pcdh8, sp5, nog* and *lhx8* were among the genes induced synergistically by Irx3-Otx2 (Supplemental Table S1). *Lhx1, chrd, gsc* and 7 other genes were synergistically induced by both combinations of transcription factors. The two groups of synergistically activated genes included ZyD H3K4me3 genes, however, they were neither enriched nor depleted significantly in these groups. These results indicated that the Foxb1-Eomes and Irx3-Otx2 combinations had cooperative roles in regulating organizer gene expression in animal caps. Together, these observations indicate that Eomes, Otx2 and the combination of Irx3 and Otx2 tend cause expression of genes that have maternally controlled H3K4me3-decoration of their promoters, whereas H3K4me3-marking of a relatively high number of genes induced in Lhx8-caps requires zygotic transcription.

## DISCUSSION

This study explores the relationships between chromatin state, regulatory elements and spatial regulation of gene expression during early development. Genome wide analysis of chromatin accessibility demonstrated pluripotent animal cap (AC) cells to have open chromatin for both ectoderm-expressed and mesoderm-expressed genes, whereas dorsal marginal zone (DMZ) cells exhibit a more restricted pattern of chromatin accessibility. This is concordant with studies that have shown that embryonic stem cells cultured *in vitro*, have a more open, accessible chromatin compared to differentiated cells (Schlesinger and Meshorer, 2019). Lineage commitment involves changes in accessibility of genes with lineage-restricted expression, many of which are accessible in pluripotent cells but inaccessible in lineages where they are not expressed. Similarly, we found that DMZ cells exhibit reduced chromatin accessibility for ectodermal genes. The earliest accessibility detected with the ATAC-seq method roughly coincides with the mid-blastula transition. This suggests that early development involves a major transition from a generally closed chromatin state to an open state that accommodates developmental competence. This is in line with studies showing that maternal transcription factors, such as Pou5f3, Sox3, Foxh1, Otx1 and Vegt, not only bind to DNA before zygotic genome activation, but also have a role in opening up chromatin (Gentsch et al., 2019; Paraiso et al., 2019).

During gastrulation, cellular heterogeneity rapidly increases beyond regional differences due to induction and morphogenesis. Analysis of blastula, early gastrula and mid-to-late gastrula single-cell data showed well-resolved cellular trajectories for non-neural ectoderm, neural ectoderm, mesoderm and organizer mesendoderm. Sequencing hand-picked single cells from dissected AC and DMZ tissue achieved both a known spatial localization and a deeper transcriptome of the cells, allowing a more detailed characterization of cell clusters. We identified organizer mesendodermal cells that express relatively high levels of Wnt inhibitors, constituting the head organizer, in addition to organizer mesodermal cells that did not abundantly express endoderm genes or Wnt inhibitors, the prospective trunk organizer. In addition, we identified cells with expression patterns consistent with superficial mesoderm and upper blastopore lip. Our analysis of whole embryo data supports the competence of *sox2/sox3*-expressing ectodermal cells for both neural as well mesodermal induction at an early gastrula stage. At the bifurcation of the neural and mesodermal trajectories, some of the cells express both *sox2* and *t (tbxt)*, which have been shown to promote respectively neural and mesodermal fates in an antagonistic fashion (Gentsch et al., 2013; Koch et al., 2017). There is a continuum between non-organizer and organizer mesoderm in the two-dimensional representation of the cells, with head and prospective trunk organizer cells exhibiting a more distinct identity.

By integrating single-cell transcriptomics and chromatin accessibility landscapes, we identified active motifs and associated transcription factors, which, when expressed in animal caps, induced organizer gene expression. *Eomes* induced *fst*, an activin and BMP antagonist, whereas *frzb*, a Wnt- antagonist, was highly enriched in *foxb1-eomes* injected caps. Eomes and other T-box transcription factors are expressed in partially overlapping expression domains, with Eomes expressed in prospective head mesoderm (Gentsch et al., 2013). Foxb1 can be induced by FGF signaling (Chung et al., 2004), and is expressed in non-involuted mesoderm and upper blastopore lip during early gastrulation (Gamse and Sive, 2001). In our data, delta-like *dll1* expression is moderately enhanced by *foxb1-eomes* overexpression in animal caps. Its expression has been reported in dorsal marginal zone and prospective mesoderm, as well as neuroectoderm (Kinoshita et al., 2011), where it is involved in lateral inhibition of neurogenesis. Whereas *foxb1* by itself induces a gene expression pattern similar to that observed in the upper blastopore lip cells, it also induced head organizer gene expression, especially in combination with *eomes*. Notably *frzb, chrd* and *gsc* are strongly induced by the Foxb1-Eomes combination. Ectopic expression of *iroquois3* in zebrafish (*iro3, irx3* in frogs) induced organizer gene expression, including that of *lhx1* and *chrd* (Kudoh and Dawid, 2001). In our experiments *irx3* by itself induced some organizer gene expression in animal caps, but in combination with *otx2* expression this was strongly enhanced. Previously we showed that the embryonic chromatin state of promoters is largely established by maternal factors (Hontelez et al., 2015). This observation extends to the promoters of genes that are activated by zygotic transcription factors. In this study we find that zygotic factors such as Fox1, Eomes, Irx3 and Otx2 predominantly act within open chromatin, using promoters that have gained the H3K4me3 permissive promoter mark by the activity of maternal factors. Interestingly the combinatorial action of the transcription factors indicates that their contributions to the early developmental program are not simply additive. We observed that the transcription factors, while strongly inducing the expression of many organizer-expressed genes, did not induce highly cluster-specific patterns of gene expression (*foxb1-eomes* C6; *irx3-otx2* C3). This indicates that their role is more selective or that their effects are further modified by other transcription factors and signaling. Single cell technologies hold great promise to analyze developmental gene regulation, especially when combined with spatial techniques. Future single cell analyses and perturbation studies will not only build on the current approaches, but will also provide the data and analytical power to reconstruct the gene regulatory networks in a spatio-temporally resolved manner.

## EXPERIMENTAL PROCEDURES

### *Xenopus* embryo manipulation

*X. tropicalis* embryos were obtained by in vitro fertilization, dejellied in 3% cysteine and collected at the desired stages. Fertilized eggs were injected with 500 ng of synthetic mRNA at the 1-cell stage and cultured until control embryos reached stage 8. Animal caps were explanted at stage 8 and cultured until stage 10.5 in 0.1x MMR. Animal use licenses were provided by DEC permits RU-DEC 2012–116, 2014–122 and CCD approval AVD1030020171826.

### Stage and tissue-specific ATAC-seq

ATAC-seq was performed as described (Bright and Veenstra, 2018). Library quality was assessed using the Agilent Bioanalyzer High-Sensitivity DNA kit for checking the fragment size distribution and signal to noise ratio was checked by performing qPCR primers spanning open and closed regions. Fragments above 700 bps were removed using AMPure XP beads to reduce unamplified clusters during sequencing. The concentration of the prepared library was quantified using Qubit and KAPA library quantification kit.

### Single cell RNA and total RNA library preparation and sequencing

Dissected embryo explants were dissociated into single cells using Ca2+ and Mg2+ free media (Sive et al., 2007). 200 cells were picked for performing single-cell RNA-seq using the modified version of Tang et al protocol (Tang et al., 2008). ERCC spike-in RNA (Thermo Fisher scientific, 4456740, 1:80000) was added to the lysis buffer. After the reverse transcription reaction, we performed 16 + 4 cycles of PCR amplification for cDNA synthesis. Amplified cDNA was purified with XP beads and the Zymo purification kit respectively and its concentration measured with Qubit® 2.0 Fluorometer (Q32866, Life Technologies). The quality of the amplified cDNA and distribution of DNA fragment size were assessed by Agilent 2100 Bioanalyzer (Agilent Technologies) with the High Sensitivity DNA Kit (5067-4626, Agilent). The sequencing libraries were prepared using the KAPA Hyper Pre-Kit (KR0961 – v6.17, Kapa Biosystems). The concentrations of the fragments with the approximate indexed adapters were quantified by KAPA library quantification. The libraries were sequenced in the Illumina platform and mapped using Bowtie (version: 1.2.2) (Langmead and Salzberg, 2012) to the indexed files of the xt9 genome.

Total RNA was extracted from animal caps using our in-house adapted Trizol-Zymo Hybrid protocol. The concentrations of all the RNA samples were measured using the DeNovix dsDNA High Sensitivity Assay (Catalog number: KIT-DSDNA-HIGH-1). The cDNA was constructed using KAPA RNA HyperPrep with RiboErase (Catalogue Number: 08278555702, Kapa Biosystems). We checked the quality of the samples by RT-qPCR using primers spanning coding regions of candidate and housekeeping genes.

### ATAC-seq alignment

The libraries were sequenced on the Illumina HiSeq 2500 with 43 bps paired-end reads each. After demultiplexing, the reads were aligned to the *X. tropicalis* genome (xt9) with bowtie2 (Langmead and Salzberg, 2012) using default settings. Reads mapping to mitochondrial DNA were excluded from the analysis together with low-quality reads including repeats and duplicates (MAPQ < 10). All mapped reads were offset by +4 bp for the +strand and −5 bp for the −strand. Peaks were called for each sample using MACS2 (Zhang et al., 2008) with parameters “-q 0.05 --nomodel --shift 37 --extsize 73”.

### ATAC-seq data analysis

Differential peaks were identified using DESeq2 (Love et al., 2014) and were used for further downstream analysis. Heatmaps were generated using fluff (Georgiou and van Heeringen, 2016). Differential peaks were then annotated to the closest gene using BEDtools (Quinlan and Hall, 2010) intersect with GREAT regions. Genome-wide boxplots of accessibility and transcriptional signal were plotted using ggplot2 (Wickham, 2016). Motif analysis on peak regions was performed using GimmeMotifs (Bruse and Heeringen, 2018).

### Single-cell RNA-seq data analysis

The dataset was filtered and quality checked for cells and genes using the package Scater (McCarthy et al., 2017). The filtered dataset was further loaded into R package Seurat (Satija et al., 2015). The hypervariable genes were used for principal component analysis, from which the statistically significant PCs were used for two-dimension t-distributed stochastic neighbor embedding (t-SNE) plots. We identified seven distinct clusters of cells using the FindClusters function in Seurat. Based on the predicted clusters, the marker genes relevant to each cluster were taken for further analysis with other datasets. Processing and visualization of the whole embryo single cell RNA-sequencing data (Briggs et al., 2018) were performed with scanpy (Wolf, Angerer and Theis, 2018). Stage 10½ AC and DMZ single cell clusters were placed in the whole embryo data based on Spearman correlations between clusters of both data sets, based on cluster mean expression of the hypervariable genes common to the two data sets.

### Integration of ATAC-seq and single cell RNA-seq

Top hypervariable genes (HVGs) from the scRNA-seq analysis were associated with the closest ATAC-seq peaks (Fig. 6A and 6B). This peak-to-gene model comprised of a matrix, with rows as peak locations and columns as expression values of target genes across the seven clusters. This was used as input of motif prediction using the gimme maelstrom function of GimmeMotifs (Bruse and Heeringen, 2018). The TFs associated with the predicted motifs were then screened based on their correlation between motif activity and their gene expression across the clusters.

## ACKNOWLEDGEMENTS

The authors are grateful to Dr. Huiqing (Jo) Zhou for helpful discussions. This work has been financially supported by the People Program (Marie Curie Actions) of the European Union’s Seventh Framework Program FP7 under REA grant agreement number 607142 (DevCom).

## Author contributions

ARB: conceptualization, investigation (performing experiments), formal analysis, visualization, writing original draft; SvG: investigation (performing experiments), visualization; QL: methodology, investigation (performing experiments); SJvH: formal analysis, software, resources; AG: investigation (performing experiments); GJCV: Conceptualization, formal analysis, funding acquisition, resources, supervision, visualization, writing original draft. All authors were involved in writing (review and editing).

## Data availability

ATAC-seq, scRNA-seq and bulk RNA-seq data have been deposited at NCBI GEO, accession number GSE145619.

## FIGURE LEGENDS

**Supplementary Figure S1**. Clustering of ATAC-seq signal at ATAC-seq peaks (center of heatmap, +/−5 kb) along with p300 and H3K4me3 ChIP-seq signals at stage 9 (blastula), 10½ (early gastrula), 12 (late gastrula) and 16 (neurula). **A.** Clustering (k 8) on all peaks. **B.** Clustering (k 6) on differential ATAC-seq peaks. H3K4me3 marked promoter regions have mostly stable accessibility across the stages, whereas differential accessibility seems restricted to enhancers (p300-bound).

**Supplementary Figure S2. A.** Genome browser view of AC and DMZ ATAC-seq at regulatory regions of key pluripotency genes (*pou5f3.3, ventx1.2, sox2* and *ventx1.1*). Most of the genes show relatively higher accessibility for animal cap cells and lower signals in DMZ. **B.** Volcano plot generated using motif activity (dot size) and differential gene expression for transcription factors identified by Gimme Motifs maelstrom analysis. The x-axis represents the log2 difference between AC and DMZ. The y-axis represents the −log10 of the adjusted p-value. (padj)). **C.** Sequence logos of few TFs having chromatin accessibility-associated motifs in AC (top row) and DMZ (bottom row).

**Supplementary figure S3.** Analysis of single cell RNA-seq data (Briggs et al., 2018). Panels depict 7705 filtered cells of stage 8, 10 and 12 embryos, shown in force-directed layout (dimensionality reduction). **A**. Cell type annotation based on marker gene expression (Briggs et al., 2018). **B**. Cell clusters (Louvain clusters L0-L20) based on hypervariable genes, labeled with predominant cell type annotation. **C.** Color-coding based on similarities in cell annotation between Louvain clusters (cf. panel B).

**Supplementary Figure S4. A.** Feature maps of marker gene expression in whole embryo scRNA-seq data. The genes were selected on the basis of published literature and for variable gene expression between single cell clusters.

**Supplementary Figure S5. A.** Heatmap showing gene expression in AC-DMZ scRNA-seq data for selected genes. **B.** Feature maps of marker gene expression in AC-DMZ scRNA-seq data. Please note that the same genes were selected for Fig. S4 and S5.

**Supplementary Figure S6.** High resolution heat map of top 100 hypervariable genes (rows) plotted for cells (columns) grouped according to cell cluster (C0-C6). Zoom in (800%) to read gene names (cf. Figure 4D).

**Supplementary Figure S7**. Whole mount *in situ* hybridization of selected genes in saggitally bisected stage 10½ embryos. Insets show lateral view of complete embryos. Arrow heads indicate the location of dorsal blastopore lip, small arrows are added at locations of expression for emphasis. Dorsal side is to the right. For *irx3*, both saggital and para-saggital bisections are shown, in addition to insets of lateral and dorsal views of whole embryo staining.

**Supplementary Figure S8**. Plots with whole embryo scRNA-seq data, showing for each cell the gene expression correlation of the cluster it belongs to, with each of the AC-DMZ clusters (C0-C6).

**Supplementary Figure S9**. High resolution heat map of differentially expressed genes (cf. Figure 6A).

