## Supplementary figures and images for "Combinatorial action of transcription factors in open chromatin contributes to early cellular heterogeneity and organizer mesendoderm specification"

### Supplemental Figure 1

Supplementary figure S1

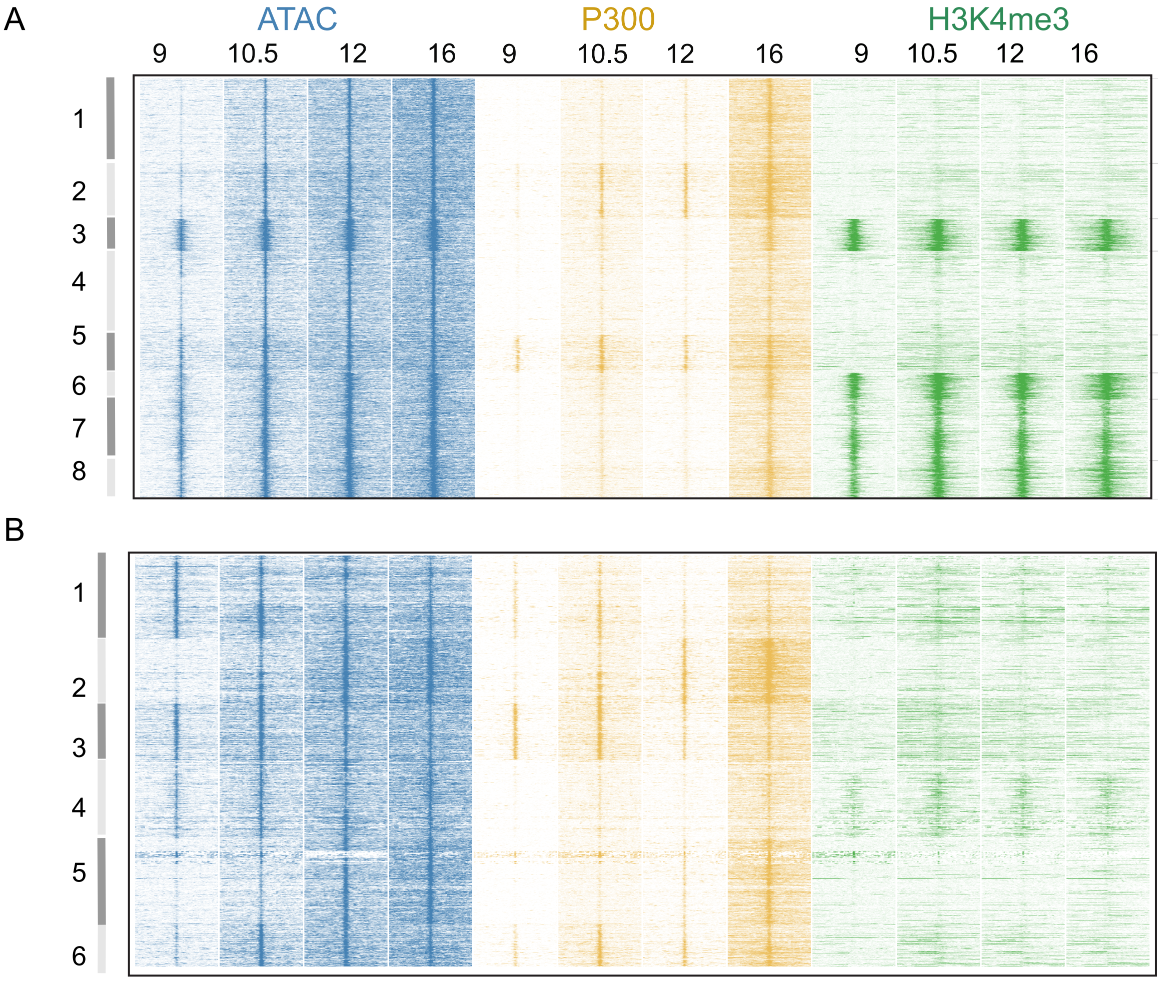

### Supplemental Figure 2

Supplementary figure S2

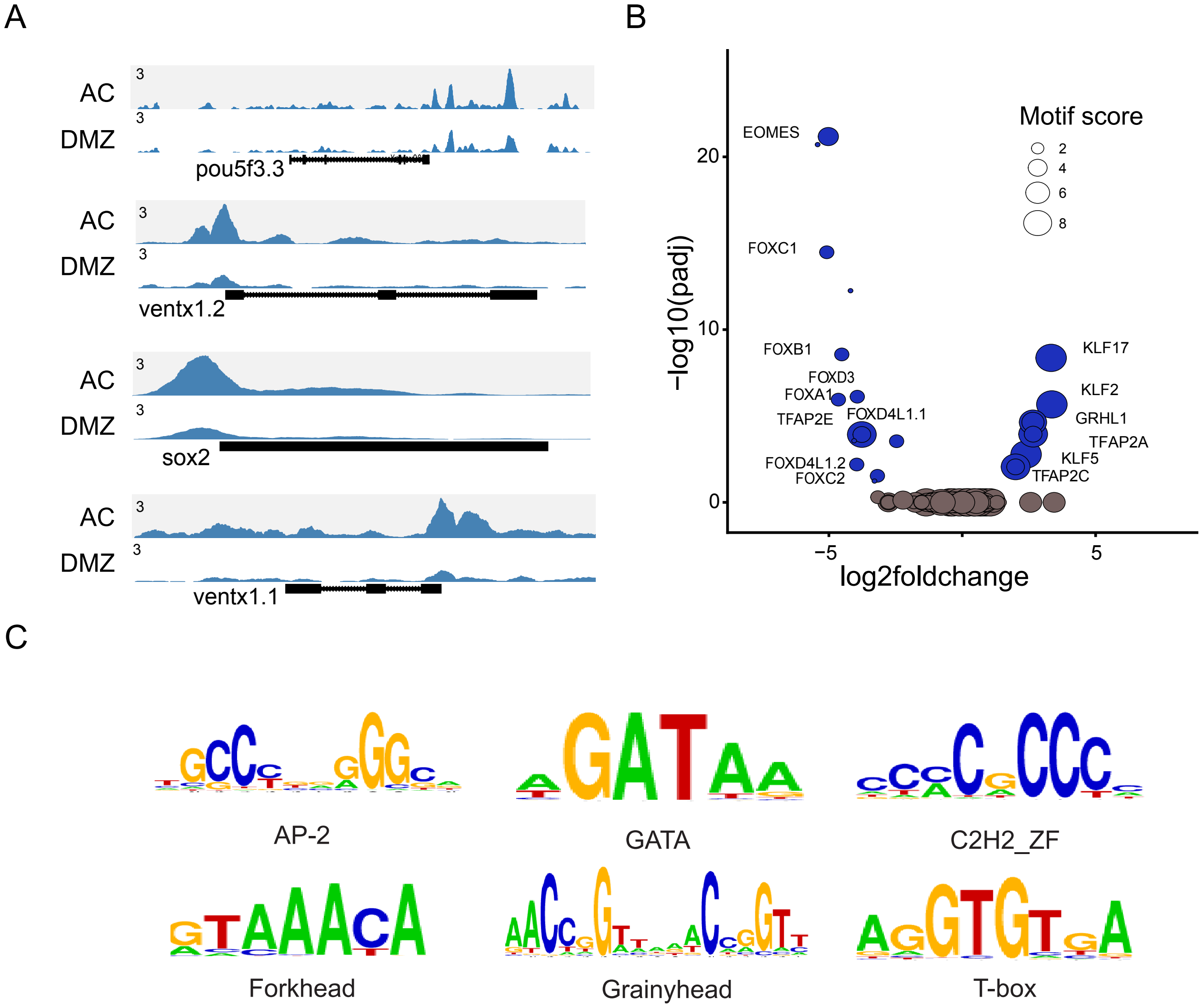

### Supplemental Figure 3

Supplementary figure S3

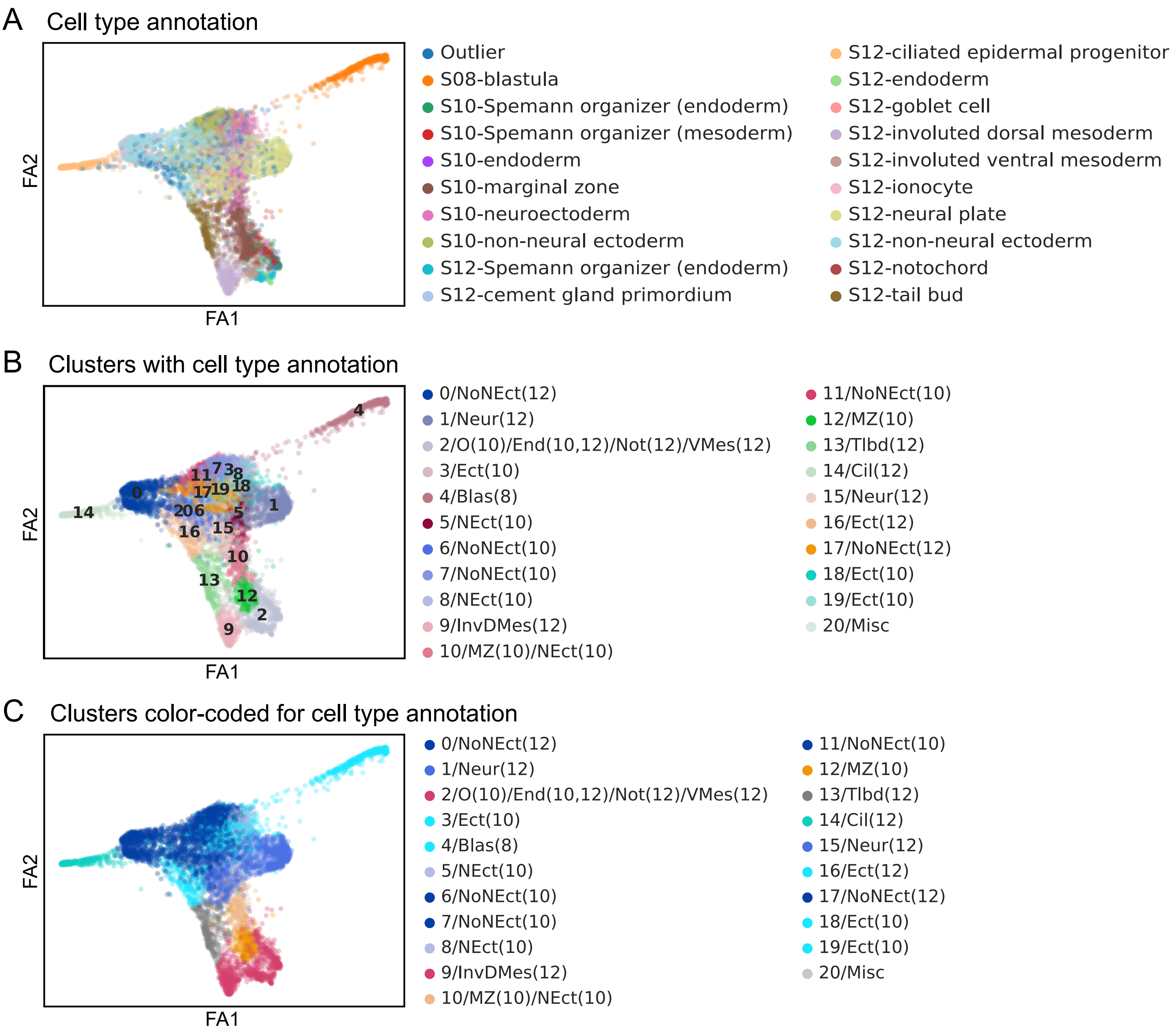

### Supplemental Figure 4

Supplementary figure S4

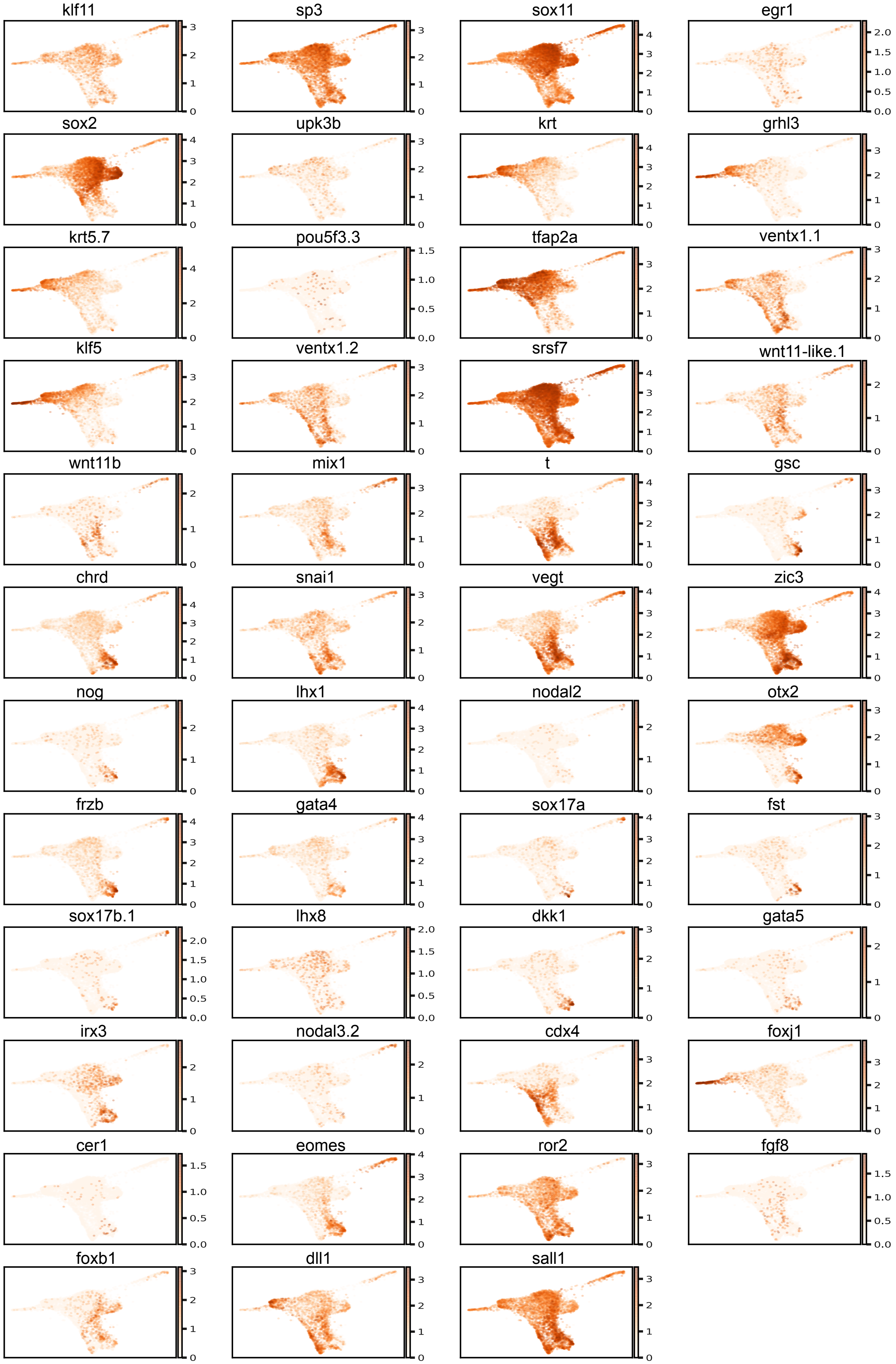

### Supplemental Figure 5

Supplementary figure 5

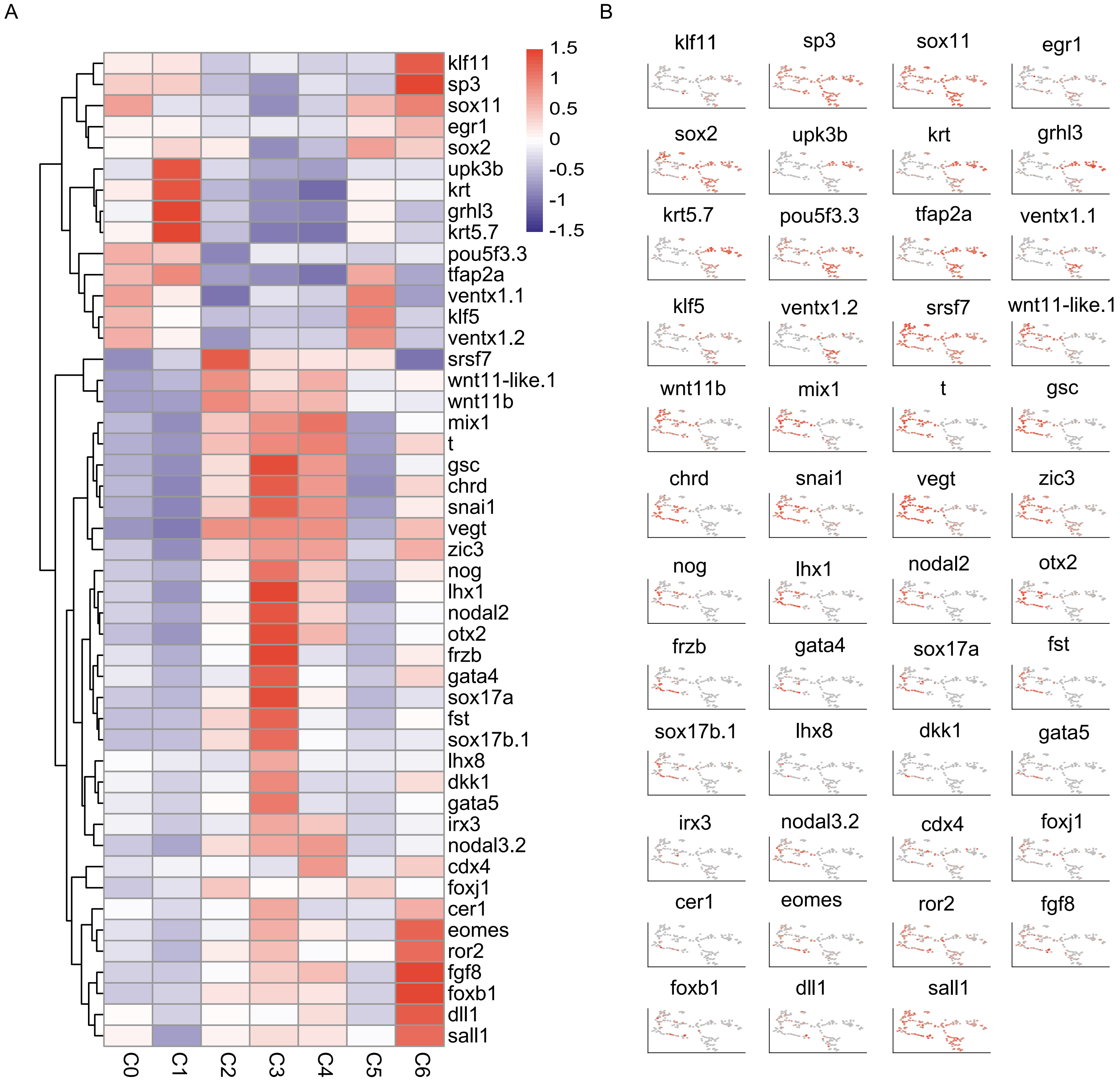

### Supplemental Figure 6

Supplementary figure S6

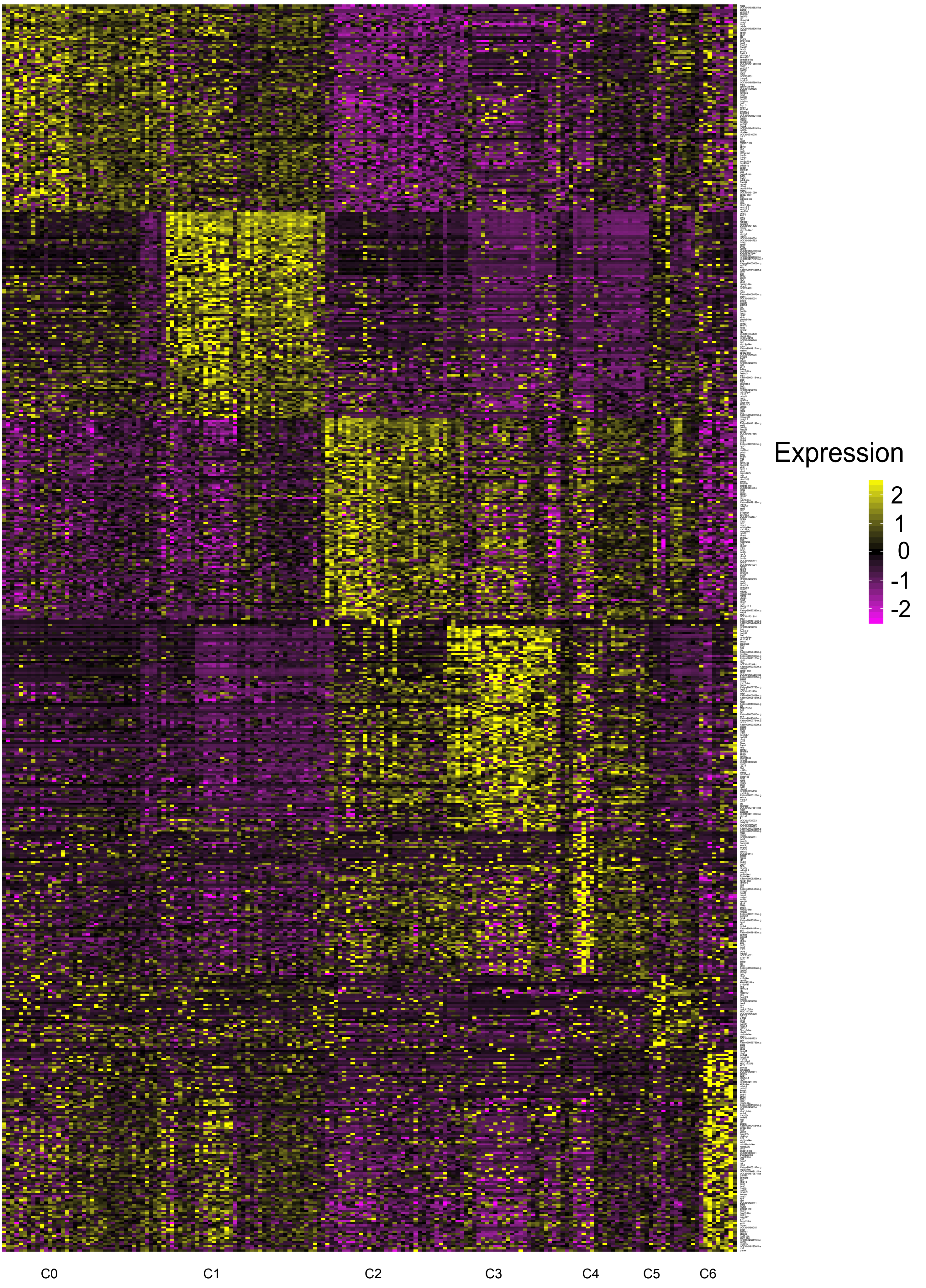

### Supplemental Figure 7

Supplementary figure S7

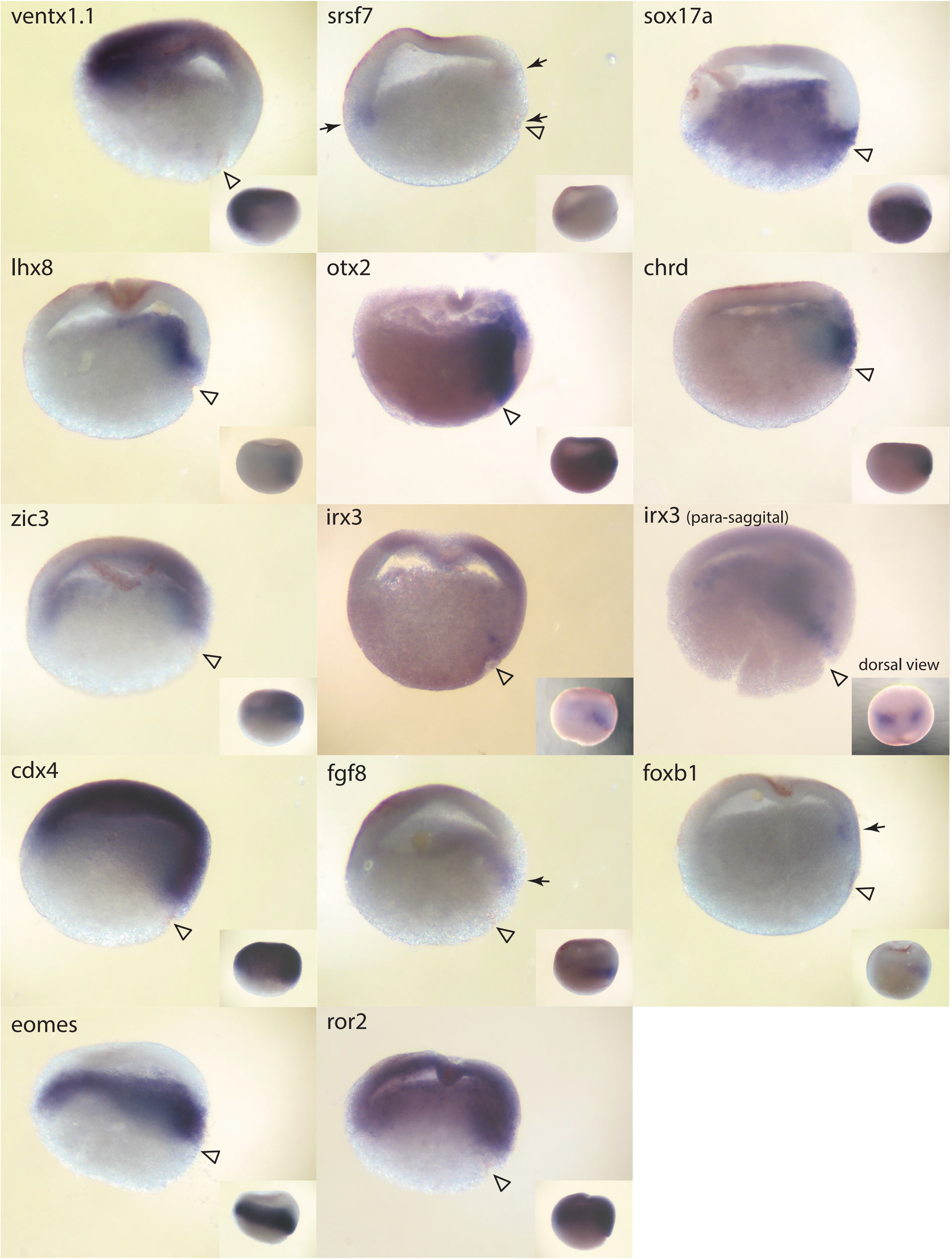

### Supplemental Figure 9

Supplementary figure S9

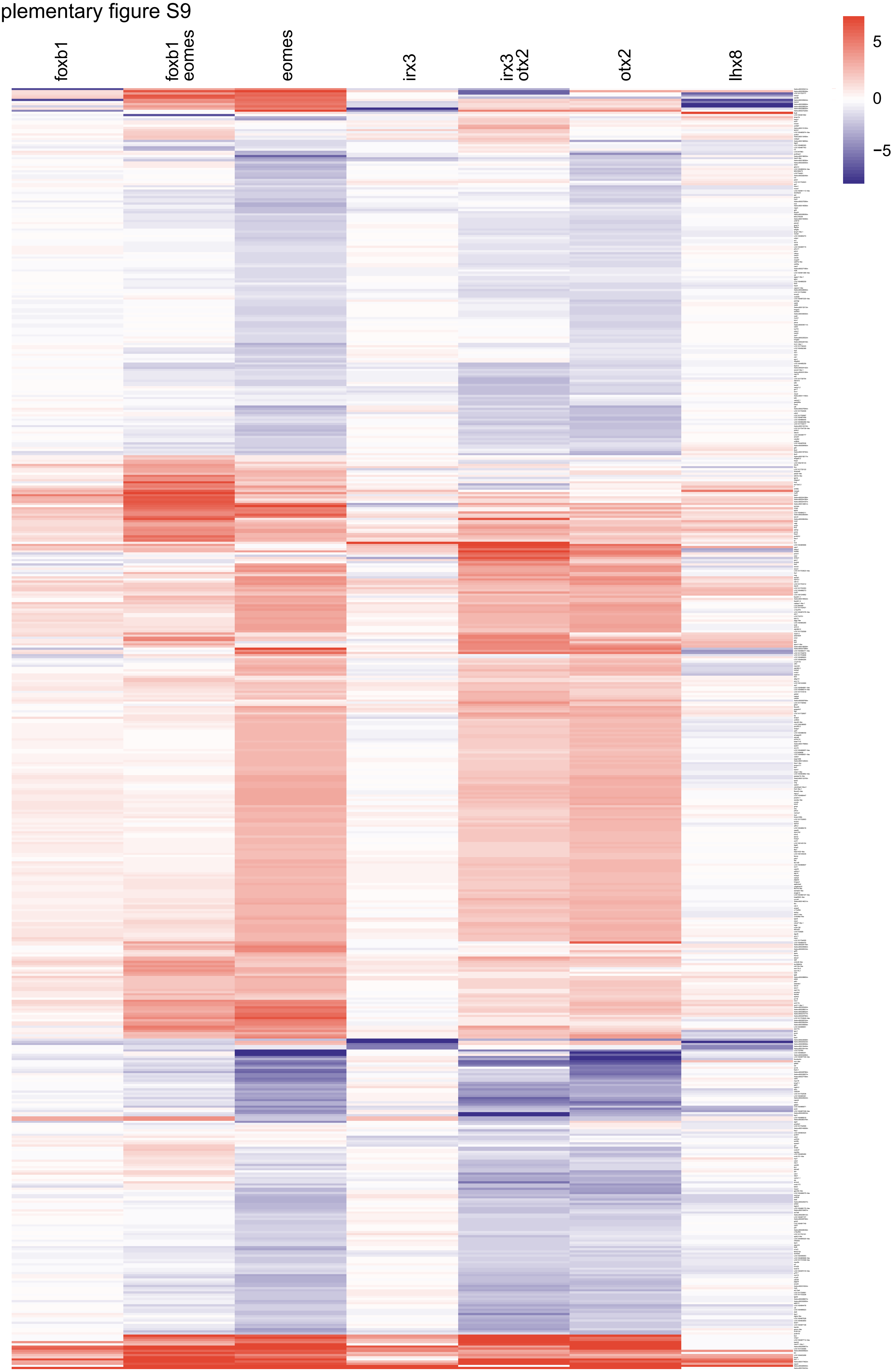
