## Supplemental Figure 8 for "Combinatorial action of transcription factors in open chromatin contributes to early cellular heterogeneity and organizer mesendoderm specification"

Supplementary figure S8

Correlation of AC and DMZ clusters C0-C6 with whole embryo stage 8, 10 and 12 clusters

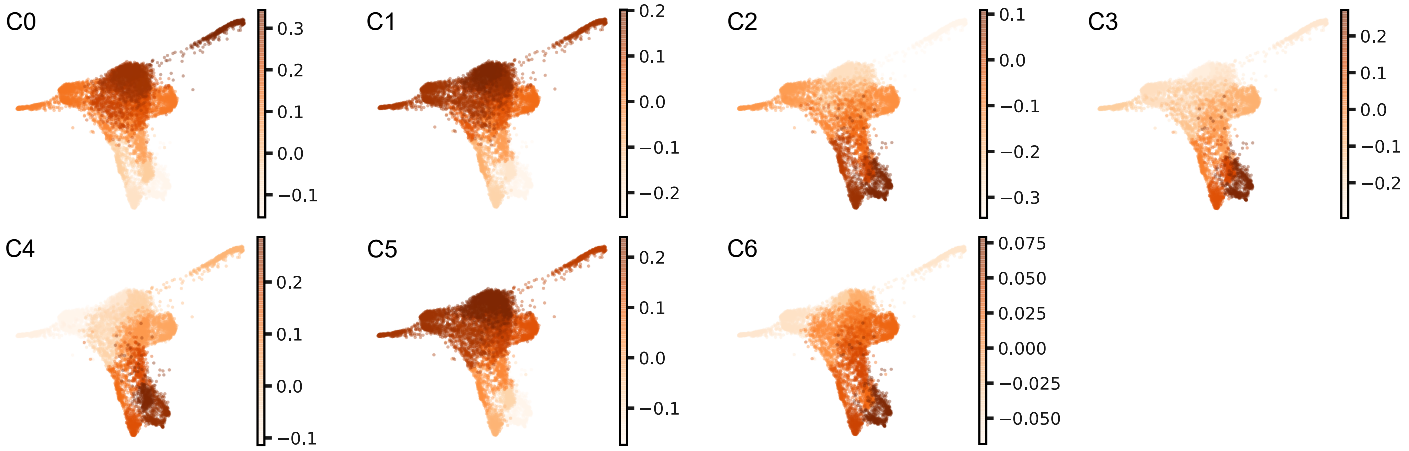
